# The Alternative Polyadenylation Factor CFIm25 Orchestrates Macrophage Antibacterial Immunity by Amplifying TAB2-Mediated MAPK and NF-κB Signaling During *Salmonella* Infection

**DOI:** 10.64898/2026.02.26.707986

**Authors:** Atish Barua, Srimoyee Mukherjee, Jeff Bourgeois, Claire L. Moore

## Abstract

Macrophage antimicrobial programs are regulated not only by transcriptional networks but also by RNA processing mechanisms affecting signal transduction and effector responses. One such mechanism, alternative polyadenylation (APA), determines mRNA fate by changing the length of the 3ʹ UTR. However, our understanding of the impact of APA on antibacterial functions and how we can manipulate it to influence infection outcomes remains limited. In this study, we identify the APA regulator CFIm25 (NUDT21) as a promoter of macrophage defense against *Salmonella enterica* serovar Typhimurium (STM). STM infection is known to drive macrophages toward an M2-like immunosuppressive state conducive to bacterial survival. Concurrent with this transition, the CFIm25 level is reduced, and the 3’ UTRs of CFIm25 targets encoding key immune proteins, such as TAB2 and TBL1XR1, are lengthened, suggesting a role for APA changes in the response to STM. Overexpression of CFIm25 in infected macrophages blocks these *Salmonella*-induced 3’ UTR changes, leading to greater mRNA and protein expression. Significantly, the increase in CFIm25 suppresses infection, thereby creating a more antimicrobial intracellular environment, improving macrophage survival, and reducing M2 properties that support bacterial replication. Specifically, CFIm25 enhances production of the antibacterial peptide LL-37, increases reactive oxygen species and nitric oxide levels, suppresses arginase activity and lactate production, and stimulates release of pro-inflammatory cytokines while inhibiting anti-inflammatory cytokines. Depletion studies show TAB2 mediates CFIm25’s antibacterial effects by activating both MAPK and NF-kB pathways. Our findings highlight APA regulation as a potential target for boosting immune defenses and developing treatments for chronic bacterial infections.

## Introduction

Macrophages play a vital role in the innate immune system by coordinating host defense through processes like phagocytosis, producing reactive oxygen and nitrogen species, and releasing inflammatory mediators (1). In a process called polarization, they can functionally evolve into either classical (M1) or alternative (M2) states, reflecting their functional diversity and plasticity. M1 macrophages combat microbes with strong antimicrobial and inflammatory responses, characterized by high levels of inducible nitric oxide synthase (iNOS), TNF-α, and IL-12 (2). In contrast, M2 macrophages support tissue repair and help resolve inflammation, expressing markers such as CD206, IL-10, and TGF-β. Balancing macrophage polarization is critical to eliminating pathogens while preventing immune-mediated tissue damage (3). Phagocytosed bacteria, such as *Salmonella enterica* serovar Typhimurium (STM), have evolved strategies to manipulate macrophage behavior, thereby creating an environment that supports their survival (4). STM’s type III secretion systems (T3SS) deliver over 40 effector proteins into host macrophages, altering signaling pathways like NF-κB and P38 MAPK, and guiding macrophages toward a non-inflammatory, M2-like state. This mechanism suppresses autophagy and reduces the production of antimicrobial mediators, thereby allowing STM to persist within macrophages for extended periods (5, 6).

While the effects of STM infection on signaling pathways and their transcriptional and protein outputs have been well studied, pathogen control of host post-transcriptional mechanisms remains poorly understood. Recent studies have highlighted alternative polyadenylation (APA) as an underappreciated post-transcriptional regulatory layer influencing immune cell phenotypes (7). APA produces mRNA isoforms with 3ʹ untranslated regions (3ʹ UTRs) of different lengths, which alter the inclusion of sequence elements that affect mRNA stability, translation, and localization (8, 9). Previously, we have found that the protein CFIm25 (NUDT21 or CPSF5), which governs 3’ UTR length by altering the poly(A) site position, is a key regulator of monocyte differentiation and macrophage polarization (10, 11). However, the role of CFIm25 in modulating bacterial infection of macrophages has not been investigate. In the current study, we show that after STM infection, the CFIm25 level is reduced, and the 3’ UTRs of CFIm25 targets encoding key immune proteins, such as TAB2 and TBL1XR1, are lengthened, suggesting a role for APA changes in the response to STM. Furthermore, restoring CFIm25 levels promotes macrophage survival and antibacterial activity during STM infection. These CFIm25-mediated effects are dependent on TAB2 and TBL1XR1. Our findings identify APA regulation by CFIm25 as an upstream modifiable mechanism that can counteract *Salmonella*-induced macrophage programming and restore macrophage bactericidal function. This study also highlights that APA is a crucial, actionable immune regulatory axis, opening new strategies to boost macrophage antibacterial responses against persistent intracellular infections.

## MATERIALS AND METHODS

### Cell Culture and Differentiation

THP-1 cells (ATCC TIB-202) were cultured in RPMI-1640 medium with 10% heat-inactivated fetal bovine serum, 2 mM L-glutamine, and 1% penicillin-streptomycin at 37°C in a humidified chamber with 5% CO₂. To induce differentiation into naive (M0) macrophages, cells were treated with three μM phorbol 12-myristate 13-acetate (PMA, Sigma-Aldrich) for 24 hours, followed by a 48-hour rest period in PMA-free medium. HEK293FT cells (Thermo Fisher; RRID:CVCL_6911) were maintained in high-glucose DMEM supplemented with 10% heat-inactivated fetal bovine serum, 2 mM L-glutamine, and 1% penicillin-streptomycin at 37°C in a humidified incubator with 5% CO₂.

### Bacterial Culture and Infection

*Salmonella enterica* serovar Typhimurium strain SL1344 was cultured overnight in LB broth at 37°C with shaking. It was then subcultured at a 1:33 ratio for 3 hours to reach log-phase growth. The bacteria were washed with PBS and resuspended in antibiotic-free RPMI-1640 medium. Prior to infection, macrophages were washed and incubated in antibiotic-free RPMI-1640 medium for 1 hour at 37°C to allow residual antibiotics to eflux and were maintained in antibiotic-free medium during the infection period. Macrophages were infected at an MOI of 1 for 30 minutes, washed three times with PBS, and then cultured in medium containing 100 μg/ml gentamicin for 1 hour to remove extracellular bacteria. Afterward, they were maintained in medium containing 25 μg/ml gentamicin for the remainder of the experiment.

### Lentivirus construction and transfection

The OE-CFIm25 lentivirus vector was a kind gift from Shervin Assassi at the University of Texas Health Science Center at Houston, TX (12). It was created by cloning the coding sequence (CDS) of human CFIm25 into the pLV-EF1a-IRES-Puro vector (Addgene), which contains an EF-1α promoter upstream of an IRES element to co-express the puromycin marker. The CFIm25 CDS was inserted between the EF-1α promoter and IRES, enabling the expression of CFIm25 and the puromycin marker from a single mRNA. The vector backbone without an insert served as the OE control.

The recombinant plasmids were co-transfected with the components of the Dharmacon™ Trans-Lentiviral packaging kit into HEK293FT cells using FuGENE® HD (Promega Corp.) according to the manufacturer’s protocol. Transfection of HEK293FT cells was performed in 6-well plates when cells were 80-85% confluent, and the transfection media were changed after 16 hours. The recombinant lentiviruses were harvested at 48 hours post-transfection, spun at 1250 rpm for 5 minutes, and filtered through a 0.45 µm filter to remove remaining cells and debris. Purified viruses were used to infect monocytic cells. 1X10^6^ target cells were seeded in 2 mL of media per well in a 6-well plate and cultured overnight. Lentiviral particles were added the next day to the cells in culture medium containing 10 µg/mL polybrene to facilitate infection. Selection of cells stably expressing OE-control and OE-CFIm25 began at 72 hours post-transfection. The growth medium was replaced with fresh selection medium containing 1 μg/mL puromycin. The puromycin-containing medium was refreshed every 2–3 days, and selection was completed after approximately 1 week. Clones were then expanded for two more weeks and then frozen for later use.

### siRNA transfection

For siRNA experiments, THP1 cells were differentiated into M0 macrophages as previously described. These M0 cells were then transfected with either gene-specific or non-targeting control siRNAs using jetOPTIMUS® transfection reagent (Polyplus), following the manufacturer’s instructions. Briefly, siRNAs were diluted in Opti-MEM and mixed with the transfection reagent for 10–20 minutes at room temperature, then added to M0 macrophages in antibiotic-free RPMI-1640 medium. Cells were exposed to the siRNA complexes for approximately 6 hours, after which the medium was replaced with complete growth medium. Subsequently, the cells were subjected to infection assays as indicated.

### ELISA

The Bio Legend® Legend Max™ kit was used to perform ELISA following the manufacturer’s protocol. In this sandwich ELISA, human TNF-α-, TGF-β-, IL-10-, or IL-12-specific monoclonal antibodies are pre-coated on a 96-well strip plate. Supernatants were collected from cells to measure TNF-α, TGF-β, IL-10, and IL-12 levels, expressed as pg/mL.

### Western blotting

Cells were lysed by adding RIPA buffer (20 mM Tris-HCl pH 7.5, 150 mM NaCl, one mM EDTA, one mM EGTA, 1% NP-40, 1% sodium deoxycholate, 2.5 mM sodium pyrophosphate, one mM β-glycerophosphate, one mM Na_3_VO_4_ and one µg/mL leupeptin) to the cell pellet, followed by incubation on ice for 15 min, centrifugation at 25,000 g for 10 min at 4°C, and supernatant collection. Protein concentration was measured with the BCA reagent (Pierce, Thermo Fisher Scientific), and 50-80 µg protein was separated on a 10% polyacrylamide-SDS gel and transferred to PVDF membrane. The membrane was then cut into segments to enable probing for multiple proteins from the same blot. The membrane was blocked for 1 hour in 5% nonfat dried milk in TBS-T (Tris-buffered saline with Tween-20; 20 mM Tris–HCl, 150 mM NaCl, pH 7.4, with 0.05% Tween-20) before overnight incubation at 4 °C with the primary antibody.

Antibodies used in this study included CFIm25 (Proteintech, 10322-1-AP), LL-37 (Santa Cruz, sc-166770), phosphorylated NF-κB P65 (Ser536) (Cell Signaling, 3033), NF-κB P65 (Cell Signaling, 8242), TAB2 (Cell Signaling, 3744), TBL1XR1 (Novus, NBP1-86996), phosphorylated STAT1 (Tyr701) (Cell Signaling, 9167), STAT1 (Cell Signaling, 14994), phosphorylated STAT3(Tyr705) (Cell Signaling, 9145), STAT3 (Cell Signaling, 4904), IκBα (Cell Signaling, 4814), phosphorylated P38(Thr180/Tyr182) (Cell Signaling, 4511), P38 (Cell Signaling, 9212), CD163 (Proteintech 68218-1-g), histone H3 (Cell Signaling, 9715), and GAPDH (Santa Cruz SC3233). Blots were developed using an enhanced chemiluminescence (ECL) substrate detection reagent (GE Healthcare) and visualized and quantified using the LI-COR Odyssey imaging systems and ImageJ software.

### Flow Cytometry

For surface marker analysis, cells were stained with fluorochrome-conjugated antibodies CD80 (BioLegend, 305208) and CD206 (BioLegend, 321110). To analyze cell death, cells were stained with propidium iodide (50 μg/ml) and evaluated for the sub-G0 population. To detect reactive oxygen species (ROS), cells were incubated with 10 μM 2’,7’-dichlorofluorescin diacetate (DCFH-DA, Sigma-Aldrich) for 30 minutes at 37°C, and ROS levels were measured by flow cytometry. For nitric oxide (NO) detection, cells were incubated with five μM 4-amino-5-methylamino-2’,7’-difluorofluorescein diacetate (DAF-FM, Invitrogen) for 30 minutes at 37°C, and NO levels were assessed by flow cytometry. Data were acquired using a BD LSRII flow cytometer and analyzed with FlowJo software (TreeStar).

### Arginase Activity Assay

Arginase activity was measured using the Arginase Activity Assay Kit (Sigma-Aldrich), which detects the conversion of arginine into urea and ornithine. The urea produced reacts with the kit’s substrate, creating a colored compound that indicates arginase activity. Briefly, cells were lysed in 100 μL of lysis buffer with protease inhibitors. After centrifugation at 10,000g for 10 minutes, the supernatant was collected, and arginase activity was determined according to the manufacturer’s instructions. The color change was read at 430 nm and normalized to total protein content.

### Lactate assay

To measure lactate released into the culture medium, conditioned media from control and STM-infected macrophage cultures were collected at 2 or 6 hours post-infection. Lactate concentrations were measured using a colorimetric lactate assay kit (Cell Biolabs MET 5012) according to the manufacturer’s instructions.

### Bacterial Colony-Forming Unit (CFU) Assay

Infected macrophages were lysed with 1% Triton X-100 in PBS. Serial dilutions were plated on LB agar and incubated at 37°C overnight. Colonies were counted and expressed as CFU per ml.

### Nuclear-Cytoplasmic Fractionation

Nuclear and cytoplasmic fractions were isolated using the NE-PER Nuclear and Cytoplasmic Extraction Reagent kit (Thermo Scientific) according to the manufacturer’s instructions.

### RT-qPCR analysis for APA and expression

RNA isolation was performed from THP-1 macrophages as previously described (10). Total RNA was extracted from 2×10^6^ cells using Trizol reagent according to the manufacturer’s instructions. 1.5 µg of RNA underwent reverse transcription with oligo dT primer and Superscript III reverse transcriptase. The resulting cDNA was amplified via qPCR with specific primers (Supplementary Figure 1B) and was quantified using the ΔΔCt method. ACTB RNA served as the normalization control for RT-qPCR-based RNA expression analyses. Primers targeting total or long transcripts were designed using the poly(A) site annotations available in poly(A)_DB, as described in detail in our previous work (14).

### Statistical Analysis

Experiments were performed with at least three independent sets. Data are presented as mean ± SE. Statistical analysis was conducted using GraphPad Prism 6.01 (GraphPad Software Inc., La Jolla, CA, USA). A two-way ANOVA was used to compare group differences. Significance levels were * = P ≤ 0.05; ** = P ≤ 0.01; *** = P ≤ 0.001. A p-value less than 0.05 was considered statistically significant (15).

## RESULTS

### *Salmonella* Typhimurium infection downregulates CFIm25 as it shifts macrophages to a replication-permissive state

A characteristic of STM infection is its ability to push macrophages toward an M2-like, replication-permissive state (16–18). This transition is reminiscent of the changes we had observed in macrophages when CFIm25 was depleted (11). To determine if CFIm25 level was altered by bacterial infection, PMA-differentiated THP-1 macrophages were infected with STM (MOI = 1). Western blotting showed rapid and significant depletion of CFIm25 protein within 6 hours post-infection (Fig. 1A). We next confirmed that the expected infection-induced changes in macrophage properties were happening in our system. The loss of CFIm25 was accompanied by decreased levels of microbicidal ROS and NO (measured by DCFH-DA and DAF-FM fluorescence respectively) (Fig. 1B, C). Another property of M2 macrophages is expression of arginase, which redirects arginine metabolism away from microbicidal NO production and toward ornithine, polyamines, and collagen precursors (19). STM infection increases arginase activity and also triggers metabolic reprogramming of macrophages to make lactate, which promotes M2 polarization and induces the SPI-2 type III secretion system (20). In our experiments, infected macrophages exhibited significantly increased arginase and lactate production (Fig. 1D, E), indicating the expected shift from an M1 to an M2 phenotype, which has been shown to support bacterial survival (21–23). These findings suggest that STM actively suppresses CFIm25 to help impair macrophage antimicrobial functions.

**Figure 1.**
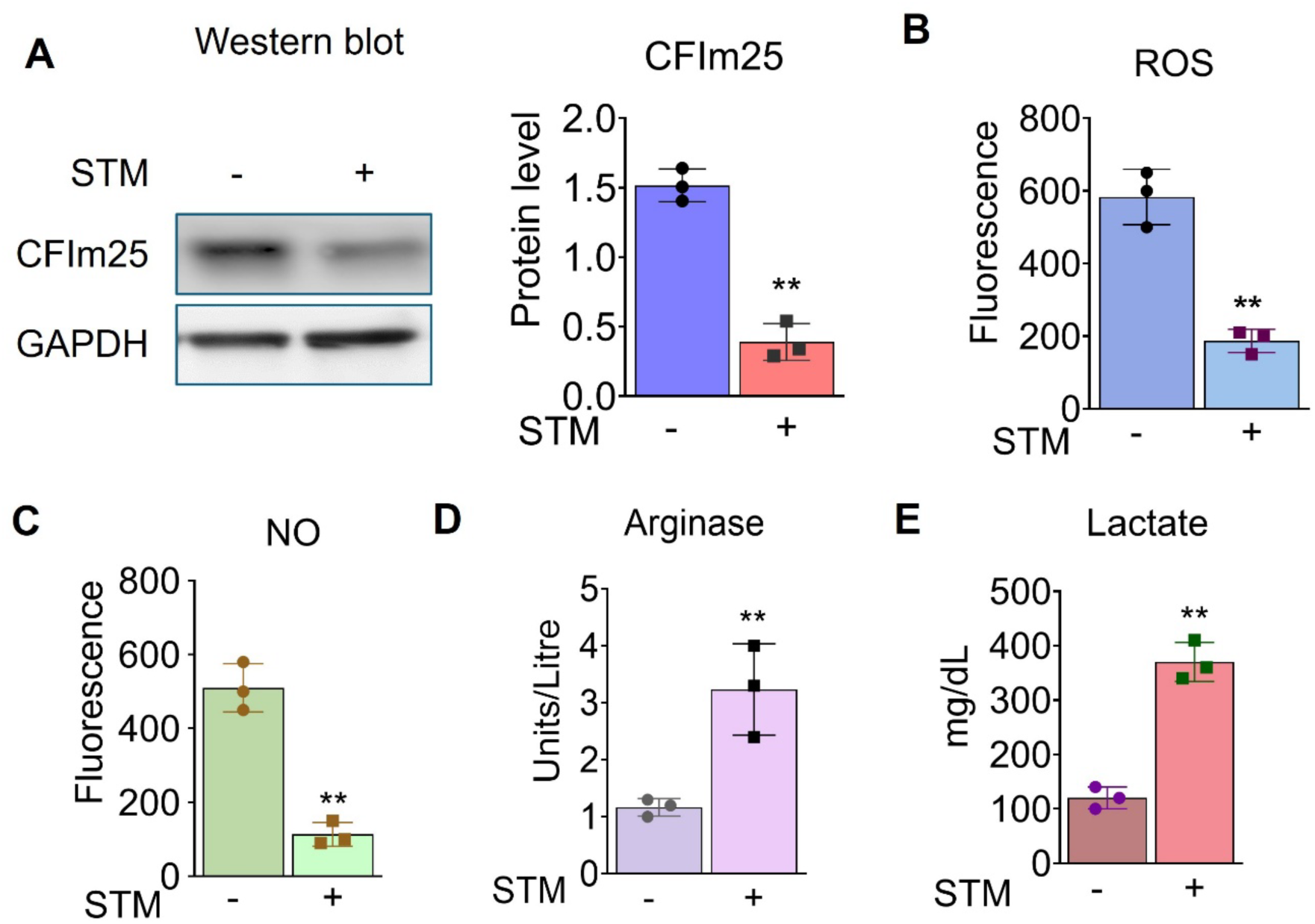
*Salmonella* Typhimurium infection downregulates CFIm25 and suppresses macrophage antibacterial activity. (A) Western blot analysis of CFIm25 protein in THP-1 macrophages six hours after infection with *Salmonella* Typhimurium (STM) compared to uninfected controls. A representative Western blot is shown on the left, and densitometric quantification of protein levels is shown on the right, normalized to GAPDH. (B-C) Flow cytometric quantification of ROS (B) and NO (C) production using DCFH-DA and DAF-FM, respectively, showing reduced microbicidal activity six hours postinfection relative to untreated macrophages. (D-E) Arginase activity, measured by urea production (D) and lactate levels (E), in macrophages six hours postinfection. All assays were compared with untreated macrophages (n = 3). Data are presented as means ± SD using three biological replicates; *, P < 0.05; **, P < 0.01.

### CFIm25 overexpression sustains the M1 phenotype and antibacterial activity during infection

To test whether CFIm25 can directly counteract the STM-driven shift from microbicidal M1 to permissive M2 macrophages, THP-1 macrophages overexpressing CFIm25 (CFIm25-OE) or control cells without CFIm25-OE were generated by lentiviral transduction and challenged with STM. Overexpression during infection was confirmed by Western blot (Fig. 2F). CFIm25-OE macrophages demonstrated superior bacterial clearance at 2 and 6 hours post-infection, with approximately 4.7 -fold (2 h) and 6.6-fold (6 h) reductions in intracellular CFU compared with control macrophages (Fig. 2A). These time points were selected to assess properties immediately after invasion (2 hours) and at the start of replication (6 hours). Moreover, the strong reduction of CFIm25 at 6 hours of infection (Fig. 1A) suggested this might be a time when progress of the infection would be susceptible to altering CFIm25 levels. To determine whether this reduction in CFU was due to macrophages killing intracellular STM or to elevated cell death in response to STM, we quantified cell death. Using flow cytometric analysis, we determined the percentage of dead cells by their ability to stain with propidium iodide and observed strong resistance to infection-induced cell death in CFIm25-OE macrophages at both 2 and 6 hours post-infection (Fig. 2B). This enhanced bactericidal activity and macrophage survival in CFIm25-OE cells correlated with increased ROS/NO output (Fig. 2C-D). In addition, conditioned medium from infected CFIm25-OE macrophages displayed increased antibacterial activity against STM, as shown by CFU assays of bacteria recovered after incubation with media from the macrophage cultures (Fig. 2E). A stable extracellular activity such as this could be due to the secretion of antimicrobial peptides by the macrophages (24). Indeed, we observed higher protein levels of the antimicrobial peptide LL-37 at both early and late time points in CFIm25 OE cells (Fig. 2F-G), indicating increased expression of this bactericidal effector.

**Figure 2.**
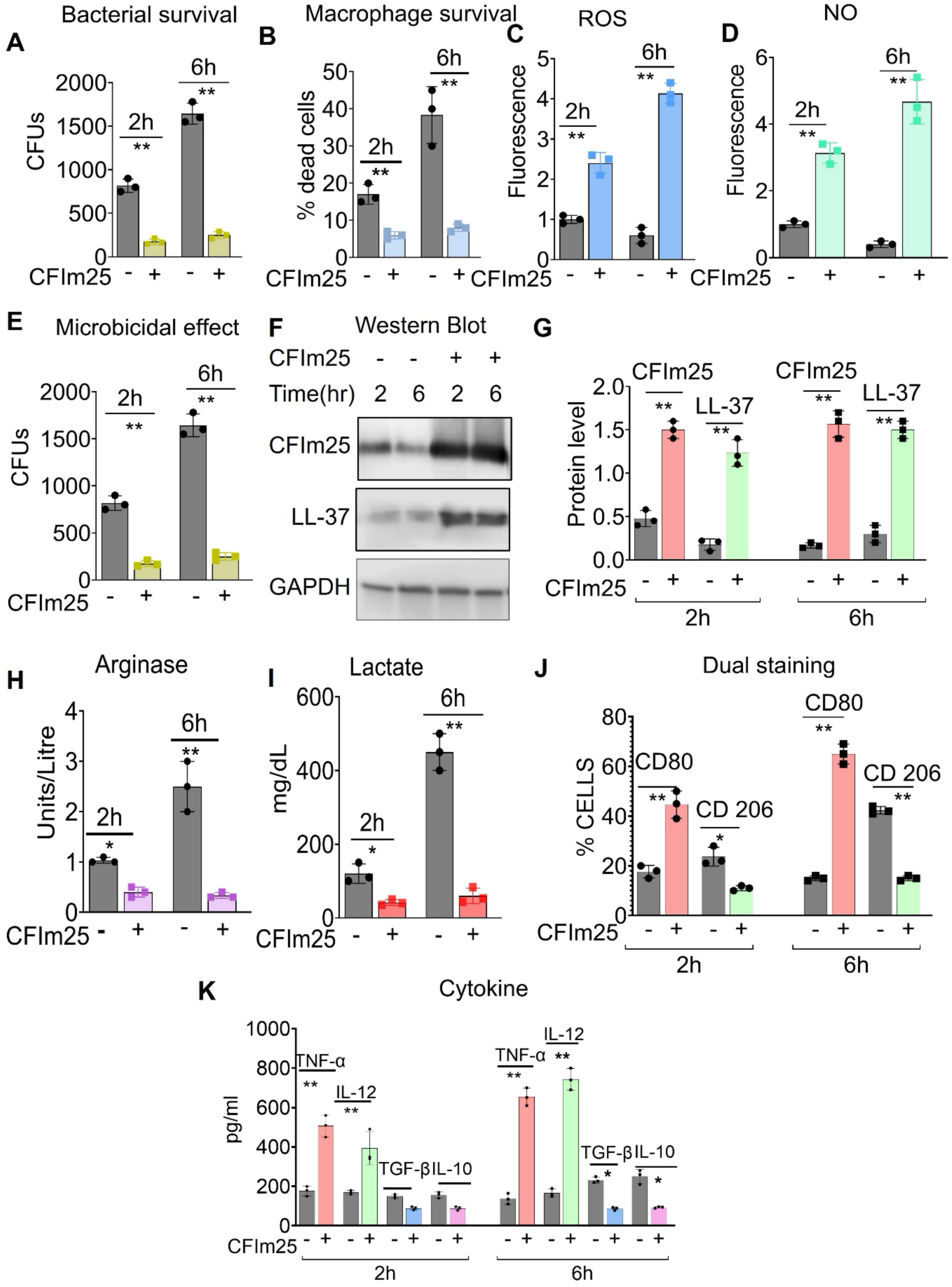
CFIm25 overexpression during STM infection enhances macrophage microbicidal function, promotes macrophage survival, and preserves M1 polarization. (A) Intracellular bacterial burden (CFU) in control (−) and CFIm25-overexpressing (CFIm25-OE, +) THP-1 macrophages at two and six hours postinfection. (B) Propidium iodide flow cytometric analysis of cell death in control and CFIm25-OE macrophages at two and six hours postinfection. Graphs show the percent of cells staining with propidium iodide; only dead cells take up the stain. (C) Quantification of ROS production using the DCFH-DA fluorescence assay. (D) Quantification of NO production using the DAF-FM fluorescence assay. (E) Antimicrobial activity of conditioned media from control and CFIm25-OE macrophages against STM, assessed by CFU recovery after incubation of bacteria with media. (F) Western blot analysis of CFIm25 and LL-37 protein levels in control and CFIm25-OE macrophages at two and six hours postinfection, (G) Quantitation of LL-37 levels by densitometry measurements and normalized to GAPDH. (H) Arginase activity measured by urea production in control and CFIm25-OE macrophages at two and six hours postinfection. (I) Lactate levels in control and CFIm25-OE macrophages at two and six hours postinfection were measured using a colorimetric lactate assay kit. (J) Flow cytometric analysis of M1 (CD80) and M2 (CD206) surface marker expression in control and CFIm25-OE macrophages at two and six hours postinfection. (K) Concentrations of the TNF-α, IL-12, TGF-β, and IL-10 cytokines in culture supernatants from control and CFIm25-OE macrophages at two h and six hours postinfection as measured by ELISA. Data are presented as means ± SD from three independent experiments (n = 3); *, P < 0.05; **, P < 0.01.

To test CFIm25’s capacity to prevent this transition to the M2 phenotype, we examined several M2 activities. We found that while controls showed a time-dependent increase in arginase and lactate levels after infection, these levels were suppressed and remained low in CFIm25 OE macrophages (Fig. 2H-I). Two additional indicators of polarization are the expression of M1- or M2-specific surface markers and the secretion of characteristic cytokines. Flow cytometric analysis showed that CFIm25-OE cells maintained high CD80 (M1) and low CD206 (M2) surface expression throughout infection (Fig. 2J). Meanwhile, controls without CFIm25 overexpression showed increased expression of the M2 marker CD206 (Fig. 2J). This shift in polarization state was confirmed by Western blot, which showed that CFIm25 suppressed expression of the CD163 M2 marker while elevating that of the HLA-DR M1 marker (Fig. 3C-D). ELISA confirmed sustained secretion of pro-inflammatory cytokines (TNF-α, IL-12) and decreased levels of anti-inflammatory cytokines (IL-10, TGF-β) in CFIm25-OE macrophages compared to controls (Fig. 2K). Overall, these findings demonstrate that increasing CFIm25 helps maintain microbicidal function during bacterial challenge by boosting ROS, NO, and LL-37, promoting macrophage survival, and preventing infection-induced M2 polarization while supporting M1-specific response features.

**Figure 3.**
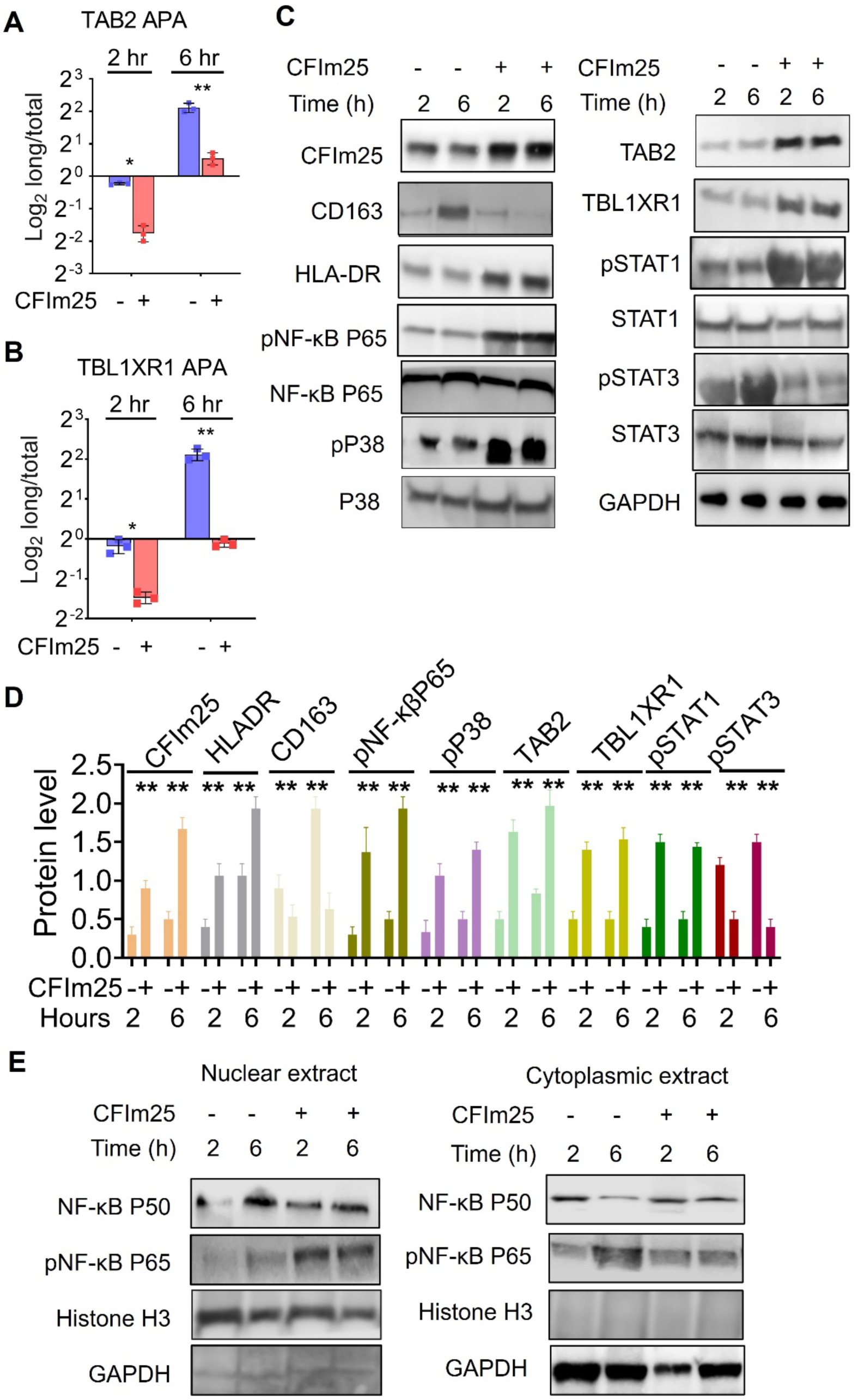
CFIm25 regulates alternative polyadenylation and the expression of key immune regulators, thereby promoting antibacterial immunity. (A-B) RT-qPCR analysis of the log2 ratio of long/total TAB2 (A) and TBL1XR1 (B) mRNA isoforms in control and CFIm25-OE macrophages at two and six hours postinfection, normalized to GAPDH. (C) Western blot analysis of CFIm25, HLA-DR, CD 163, and the indicated immune signaling proteins (“P”, protein; “p”, phosphorylated form) in control and CFIm25-OE macrophages at two and six hours postinfection, with GAPDH as the loading control. (D) Quantification of changes of the indicated proteins in Western blots of samples from control and CFIm25-OE macrophages at two and six hours postinfection, with values normalized to GAPDH. (E) Representative Western blot showing nuclear and cytoplasmic distribution of NF-κB subunits in control and CFIm25-OE macrophages two and six hours postinfection, assessed by fractionation and immunoblotting. Data from all quantifications are presented as means ± SD from three independent experiments. *, P < 0.05; **, P < 0.01.

### CFIm25 modulates APA of key components of the NF-κB pathway and affects multiple signaling pathways governing M1/M2 fate

Our previous study showed that CFIm25 OE in monocytes expedites macrophage differentiation (10). It activates the NF-κβ signaling pathway, causes shortening of the 3’ UTRs of TAB2 and TBL1XR1 transcripts, and increases the protein levels of these NF-κβ positive effectors (10). We then aimed to assess how STM infection influenced APA of these targets and performed RT-qPCR with primer pairs that detected either total mRNA levels or specifically those with longer 3’ UTRs (Supplementary Fig. 1B-C). This analysis revealed that STM infection caused an extension of the 3’ UTRs of TAB2 and TBL1XR1 mRNAs in control cells, indicating increased use of distal polyadenylation sites (Fig. 3A). However, CFIm25-OE cells maintained a lower long/total mRNA ratio during infection, signifying greater use of proximal sites and 3’ UTR shortening (Fig. 3A-B). This shortening was associated with significantly increased transcript and protein levels of both TAB2 and TBL1XR1 (Fig. 3C-D and Supplementary Fig. 1A).

We next examined activation of signaling pathways that regulate M1 and M2 phenotypes of specific kinases. Infected CFIm25 OE macrophages also exhibited increased phosphorylation of STAT1, NF-κB P65, and P38 MAPK, which drives M1 polarization (26–28). A hallmark of STM infection is the activation of STAT3, which leads to increased expression of anti-inflammatory genes, such as IL-10 and IL-4Ra, and promotes the M2 state. (29–31). Consistent with CFIm25’s ability to resist the STM-induced push to M2-like cells, CFIm25 OE blocked STAT3 phosphorylation during infection (Fig. 3C-D).

An effective activation of NF-κB-dependent genes requires the translocation of phosphorylated NF-κB P65 into the nucleus, increased association of P65 with the NF-κB P50 subunit, and a decrease in repressive nuclear-localized P50-P50 homodimers (32, 33). To assess the effects of CFIm25 on localization, we conducted nuclear-cytoplasmic fractionation on infected cells with and without CFIm25 overexpression. We confirmed a clean fractionation by the absence of GAPDH in the nuclear fraction and absence of histone H3 in the cytoplasmic one (Fig. 3E). This experiment showed that CFIm25-OE macrophages had markedly higher nuclear phosphorylated p65 at two and six hours post-infection compared with controls. In contrast, the control cells showed very little nuclear phosphorylated P65 but increased nuclear NF-κB P50 at 6 hours (Fig. 3E). This indicates a shift toward repressive P50-P50 homodimers during infection, which CFIm25 can reverse by promoting activating P65-containing heterodimers. Overall, these data suggest that sustained CFIm25 activity protects macrophages against STM-induced immunosuppressive reprogramming by maintaining a signaling environment that favors M1-like inflammatory polarization.

### TAB2 and TBL1XR1 are critical for CFIm25-mediated antimicrobial defense

We previously showed that knockdown of TAB2 and TBL1XR1 in CFIm25-overexpressing macrophages attenuated CFIm25-mediated NF-κB activation. We now wanted to determine if depletion of TAB2 or TBL1XR1 would block the protective effects of CFIm25 during infection. CFIm25-OE macrophages were transfected with control or siRNAs against TAB2 or TBL1XR1, followed by STM infection for 6 hours and harvesting of cells. The knockdown caused a pronounced depletion of both proteins (Fig. 4H). TAB2 knockdown produced a striking increase in the amount of recovered bacteria while TBL1XR1 knockdown caused a more modest increase (Fig. 4A), indicating a reduction in CFIm25-mediated bacterial clearance. TAB2 and TBL1XR1 siRNAs also suppressed TNF-α and IL-12 secretion, as well as ROS and NO levels, with TAB2 siRNA having a stronger effect (Fig. 4B-D). Both knockdowns comparably and strongly reduced LL-37 levels (Fig. 4H). TAB2 knockdown upregulated the anti-inflammatory cytokines TGF-β and IL-10, arginase activity, and lactate production, whereas TBL1XR1 had no or only a modest effect on these M2 properties (Fig. 4B, E, and F). Flow cytometric analysis indicated that TAB2 depletion reduced the M1 marker CD80 and increased expression of the M2 marker CD206 (Fig. 4G).

**Figure 4.**
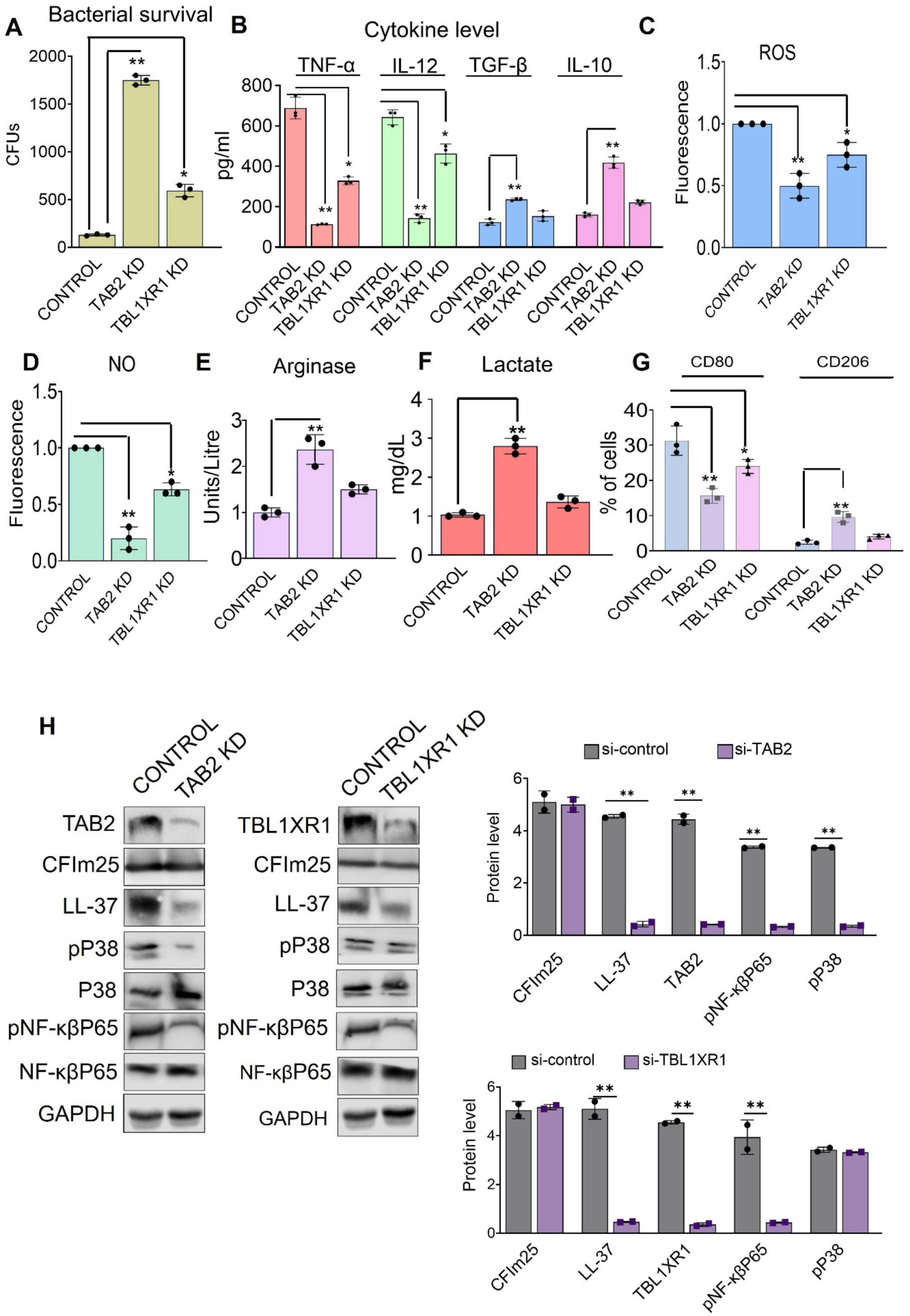
TAB2 and TBL1XR1 are critical for CFIm25-mediated antimicrobial defense. (A) CFUs recovered at six hours postinfection from Intracellular STM in CFIm25-OE macrophages transfected with control siRNA, TAB2 siRNA, or TBL1XR1 siRNA. (B) Concentrations of TNF-α, IL-12, TGF-β, and IL-10 in culture supernatants from CFIm25-OE macrophages expressing control, TAB2, or TBL1XR1 siRNAs, harvested at six hours postinfection, and measured by ELISA. (C-F) ROS (C), NO (D), arginase activity (E), and lactate (F) levels at six hours postinfection in CFIm25-OE macrophages treated with control, TAB2, or TBL1XR1 siRNA. (G) Flow cytometric analysis of CD80 and CD206 expression at six hours postinfection in CFIm25-OE macrophages treated with control, TAB2, or TBL1XR1 siRNA. (H) Western blot analysis of the indicated NF-κB and MAPK signaling proteins at six hours postinfection in CFIm25-OE macrophages treated with control, TAB2, or TBL1XR1 siRNA, with densitometric quantification from three independent experiments (“P”, protein; “p”, phosphorylated form). Data are presented as means ± SD (n = 3); *, P < 0.05; **, P < 0.01.

NF-κB P65 and P38 MAPK activation both enhance anti-*Salmonella* immunity by inducing proinflammatory, M1-like macrophage responses. Conversely, *Salmonella* often inhibits these pathways to escape immune detection. (34, 35). When levels of phosphorylated P65 and P38 remain elevated, macrophages increase the production of antimicrobial agents, including ROS, NO, LL-37, and inflammatory cytokines, thereby helping to control intracellular bacterial growth and survival. As expected from their known roles in NF-κB signaling, both TAB2 and TBL1XR1 depletions reduced the level of phosphorylated P65 (Fig. 4H). Consistent with TAB2’s role in activating the P38 MAPK pathway (36, 37), silencing TAB2, but not TBL1XR1, decreased P38 phosphorylation (Fig. 4H), indicating a central role for TAB2 in integrating NF-κB and MAPK signaling for antibacterial immunity.

## Discussion

This study highlights the importance of CFIm25, an APA regulator, in determining whether STM– infected cells become bactericidal or permit bacterial replication. Upon STM infection, CFIm25 is rapidly downregulated, thereby lengthening the 3ʹ UTRs of mRNAs encoding the critical signaling proteins TAB2 and TBL1XR1. This change is associated with reduced ROS and NO production, and a metabolic shift toward arginase- and lactate-rich, M2-like macrophages that facilitate bacterial survival. Conversely, maintaining CFIm25 expression reverses this shift at both APA and multiple functional levels. It promotes proximal poly(A) site usage in TAB2 and TBL1XR1 transcripts and an increase in their encoded proteins, enhances NF-κB P65- and P38-driven pro-inflammatory responses, and preserves an M1-like macrophage phenotype evident from upregulation of M1 markers and production of ROS, NO, the LL-37 anti-microbial peptide, and pro-inflammatory cytokines, all leading to enhanced intracellular STM elimination and improved macrophage survival.

Large-scale APA datasets now provide a framework for interpreting these mechanistic observations. Quantitative 3ʹ UTR-APA maps across human immune cell types reveal thousands of immune response–associated 3ʹ UTR APA quantitative trait loci (3ʹaQTLs), which link germline genetic variation to cell type–specific shifts in proximal versus distal poly(A) site usage at key immune genes (38, 39). Many of these 3ʹaQTLs are dynamically unmasked only after stimulation, underscoring that APA is a major, stimulus-responsive regulatory layer in innate and adaptive immunity. Single-cell 3ʹ-end profiling across hematopoietic and lymphoid lineages further demonstrates lineage- and state-specific APA programs, with systematic 3ʹUTR shortening of immune effector and signaling genes accompanying activation and differentiation (38, 40).

Consistent with the 3ʹaQTL analysis, global 3ʹ UTR shortening is observed during the innate immune response to bacterial infections, including *Listeria monocytogenes* and *Salmonella* Typhimurium. (41) and during infection of macrophages with vesicular stomatitis virus (42), indicating that infection-driven modulation of 3ʹ end processing can broadly rewire innate defenses. The immune 3ʹaQTL atlases show that many APA-sensitive loci encode upstream regulators and negative-feedback components of the NF-κB and MAPK pathways, suggesting that 3ʹ UTR selection determines how strongly and for how long these pathways remain active after stimulation. Within this broader landscape, our data place CFIm25 as a key macrophage 3ʹ UTR-APA regulator whose manipulation is sufficient to override *Salmonella*’s attempt to impose an M2-like, replication-permissive program and instead switch macrophages to a bactericidal state.

Mechanistically, our data support a model in which CFIm25-driven 3ʹ UTR shortening of TAB2 strengthens the TAK1–TAB signalosome, thereby enhancing NF-κB activation and P38 signaling in infected macrophages. (43). The CFIm25-mediated increase in TBL1XR1 would also contribute by facilitating the nuclear localization of NF-κB P65 (44, 45). During STM infection in CFIm25-overexpressing cells, this amplification is accompanied by increased nuclear NF-κB p65, reduced nuclear NF-κB p50 homodimers, and a shift toward higher STAT1 and lower STAT3 phosphorylation, a transcription factor balance tightly linked to M1 polarization and antimicrobial effector gene induction (46). Conversely, downregulation of CFIm25 by STM correlated with reduction of TAB2 and TBL1XR1 expression and weakening of the NF-κB/P38 output. CFIm25-mediated regulation of TAB2 and TBL1XR1 levels likely combines with STM*-*driven STAT3 activation to enforce an M2-like, infection-permissive phenotype (47, 48). Interestingly, a recent study has reported that *Mycobacterium tuberculosis* also targets CFIm25 but through a different mechanism. Unlike STM, which reduces CFIm25 levels, this pathogen supports its survival in macrophages by disrupting CFIm25’s interaction with another subunit of the CFIm complex (49).

Taken together, these results and the broader APA datasets suggest that 3ʹ UTR-APA acts as a post-transcriptional “decision layer” that integrates multiple cues to direct macrophage polarization toward either bacterial clearance or tolerance. These findings imply that manipulating APA regulators, such as CFIm25, could open new host-targeted therapeutic avenues to reprogram infected macrophages in chronic bacterial infections.

---

**Supplementary Figure S1.**
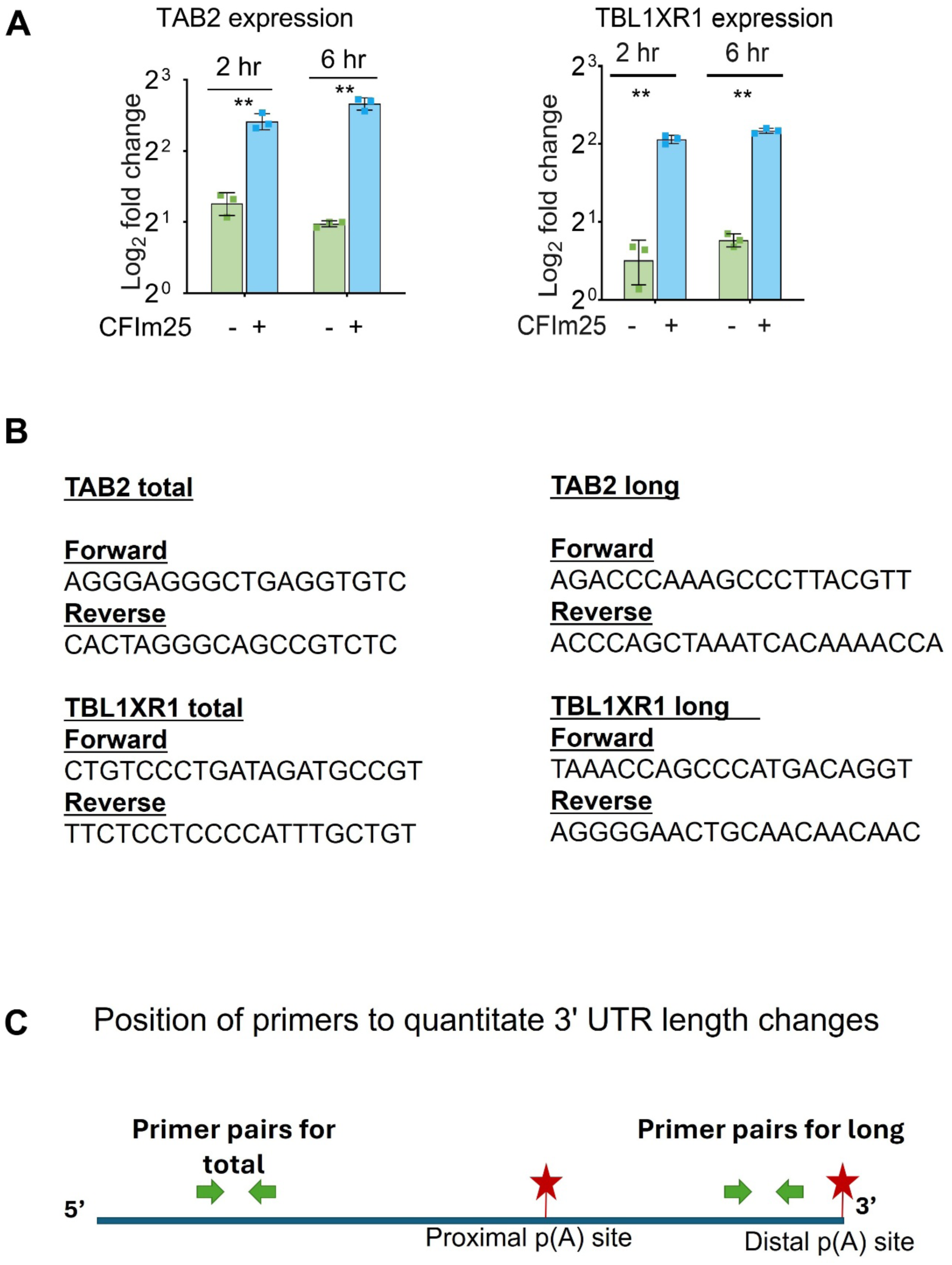
(A) CFIm25 overexpression increases TAB2 and TBL1XR1 mRNA expression after STM infection, assessed by RT-qPCR analysis in control and CFIm25-OE macrophages at 2 and 6 hours postinfection with STM (n = 3). Data are presented as means ± SD (n = 3); *, P < 0.05; **, P < 0.01. (B) Sequence of primers used for RT-qPCR analysis of TAB2 and TBL1XR1 APA. (C) Schematic representation of the long/total TAB2 and TBL1XR1 transcript regions targeted for RT-qPCR to assess APA and mRNA expression.

## Authors’ contributions

CM and AB conceived the study. CM provided general oversight. AB, JB, and SM developed the strategy and methodology, designed the experiments, acquired the data, reported the results, and organized the findings. All authors interpreted the results. AB wrote the first draft of the manuscript, and all authors commented on subsequent versions of the manuscript. All authors contributed to the article and approved the submitted version.

## Acknowledgements

We want to acknowledge the Tufts University Flow Cytometry Core and its staff for their support with flow cytometry data acquisition and analysis. Their expertise and resources were invaluable to this study. We are also grateful to Dr. Linden Hu, Department of Molecular Biology and Microbiology, Tufts School of Medicine, for his guidance in helping us get started on this project.

